# Automating multimodal microscopy with NanoJ-Fluidics

**DOI:** 10.1101/320416

**Authors:** Pedro Almada, Pedro M. Pereira, Siân Culley, Ghislaine Caillol, Fanny Boroni-Rueda, Christina L. Dix, Romain F. Laine, Guillaume Charras, Buzz Baum, Christophe Leterrier, Ricardo Henriques

## Abstract

Fluorescence microscopy can reveal all aspects of cellular mechanisms, from molecular details to dynamics, thanks to approaches such as super-resolution and live-cell imaging. Each of its modalities requires specific sample preparation and imaging conditions to obtain high-quality, artefact-free images, ultimately providing complementary information. Combining and multiplexing microscopy approaches is crucial to understand cellular events, but requires elaborate workflows involving multiple sample preparation steps. We present a robust fluidics approach to automate complex sequences of treatment, labelling and imaging of live and fixed cells. Our open-source NanoJ-Fluidics system is based on low-cost LEGO hardware controlled by ImageJ-based software and can be directly adapted to any microscope, providing easy-to-implement high-content, multimodal imaging with high reproducibility. We demonstrate its capacity to carry out complex sequences of experiments such as super-resolved live-to-fixed imaging to study actin dynamics; highly-multiplexed STORM and DNA-PAINT acquisitions of multiple targets; and event-driven fixation microscopy to study the role of adhesion contacts in mitosis.

## Introduction

Fluorescence microscopy is ubiquitously used to observe cellular processes, thanks to its ease of use, exquisite sensitivity and molecular specificity. It is generally performed using dedicated sample preparation procedures, tailored to achieve optimal imaging conditions for each chosen technique. Obtaining the best possible temporal and spatial resolution, while keeping a high signal, capacity for deep imaging and live-cell compatibility is nearly impossible with a “one-size fits all” approach. As such, each microscopy method entails a compromise between some of these features (1). Alternatively, unique insights can be gained by combining information from multiple approaches, but at the cost of complex correlative workflows (2, 3). Recent developments toward high-resolution imaging of a large number of molecular targets have further broadened the variety and complexity of imaging procedures, with the use of multiple rounds of labelling and imaging (4–7). These elaborate protocols involve sequences of sample imaging, washing and labelling. Carrying these protocols in a non-automated manner critically hampers their reproducibility and throughput, limiting their appeal for quantitative work (8).

Automated fluid handling using microfluidic chips presents an attractive alternative, but adds specific constrains on culturing conditions and sample preparation (5, 9). A more simple and tractable method would automate fluid exchange in commonly used open imaging chambers, in a manner similar to standard laboratory protocols, while being easily adaptable to existing microscope stages. For this, we devised an easy-to-implement open-source system called NanoJ-Fluidics (Fig. 1a-b), which primarily consists of an automated computer-controlled syringe pump array capable of rapidly and robustly exchanging sample conditions. This allows the automation of the protocols of sample treatment, labelling and preparation directly on the microscope stage (Fig. 1c and S1). Our approach makes complex multimodal imaging protocols highly accessible to researchers. To demonstrate this, we performed a range of complex multistep experiments: first, we devised live-to-fixed experiments with time-lapse imaging of living cells expressing reporter probes, followed by *in-situ* online fixation, labelling and super-resolved imaging, thus connecting the dynamics and structural dimensions of cellular processes (Fig. 2 and Movie S1). Next, we used NanoJ-Fluidics to automate a highly-multiplexed STORM and DNA-PAINT acquisition to obtain a five-colour nanoscopic image of distinct cellular targets (Fig. 3 and Movie S2). Finally, we showcased the capacity for high-content correlative imaging with event-driven fixation, studying the role of adhesion contacts for cells in mitosis (Fig. 4 and Movie S3).

**Fig. 1.**
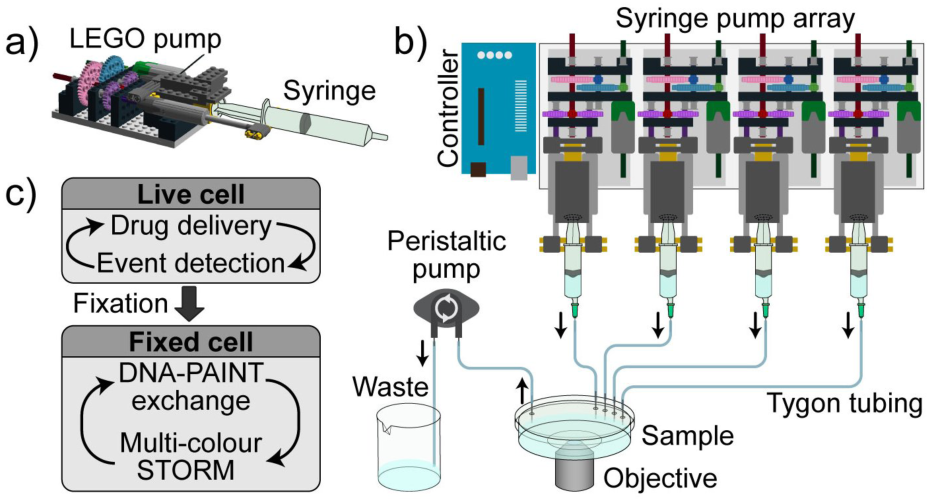
Schematics of the NanoJ-Fluidics system. a) 3D side view of a single LEGO syringe pump unit with syringe attached. b) 2D top view of a syringe pump array (representing 4 pumps out of 128 maximum) and a fluid extraction peristaltic pump, both controlled by an Arduino^®^ UNO. c) Example of possible workflow.

**Fig. 2.**
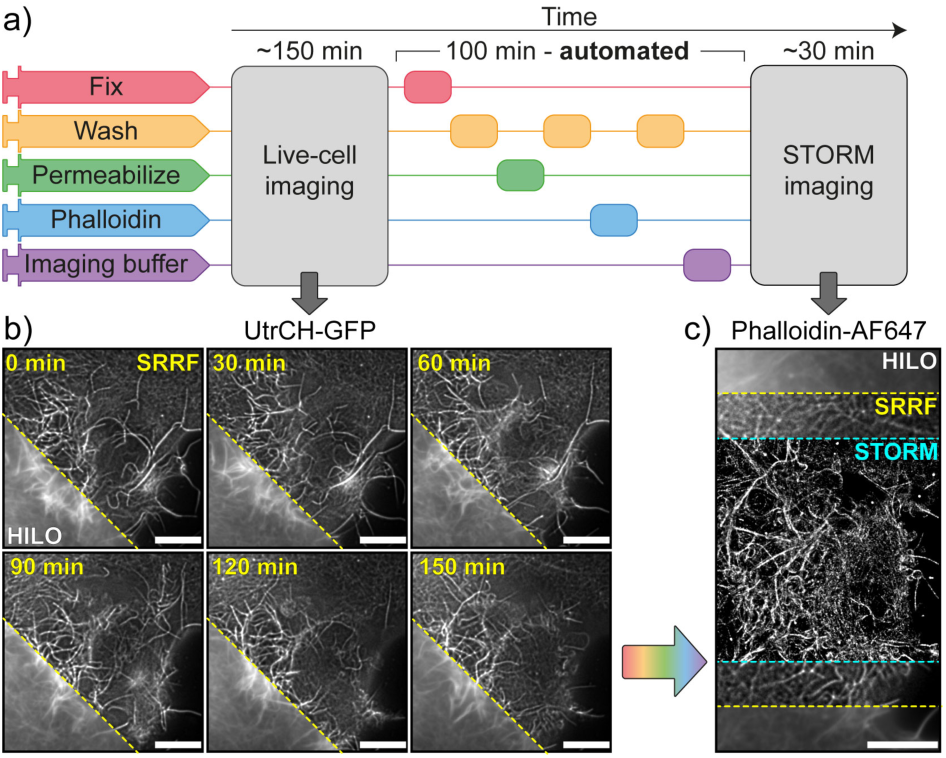
Super-resolution live-to-fixed cell imaging of actin in COS-7 cells. a) NanoJ-Fluidics workflow used for live-to-fixed super-resolution imaging. b) HILO and SRRF microscopy images of a COS-7 cell expressing UtrCH-GFP imaged every 10 min for 150 min (a zoomed region of the imaged cell is shown at 30 minutes intervals, Sup. Movie S1 shows extended time-lapse and field-of-view). c) HILO and SRRF microscopy images of UtrCH-GFP at t = 150 min and the corresponding STORM image after fixation and staining with phalloidin-AF647. Scale bars are 10 μm.

**Fig. 3.**
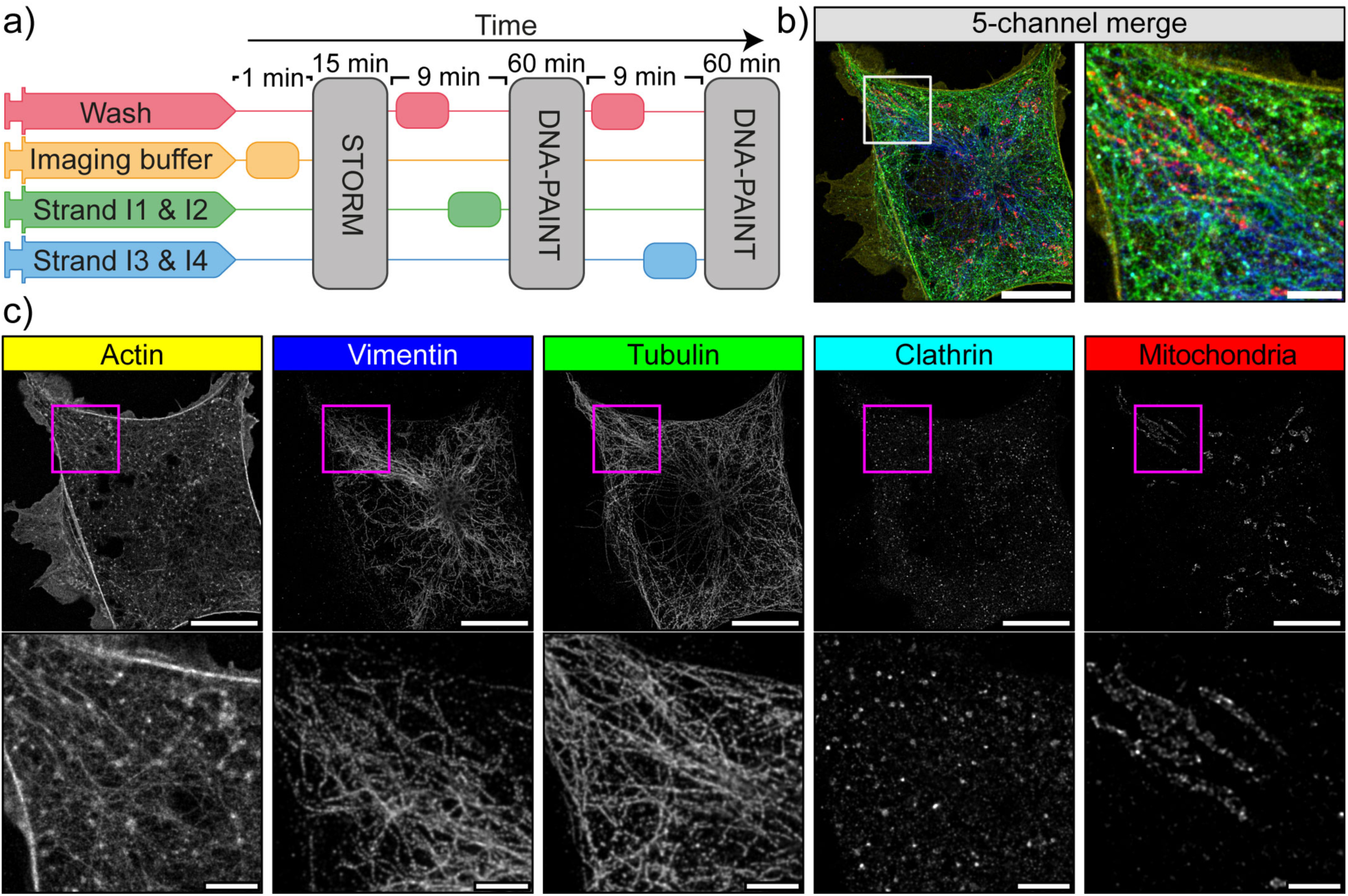
**Automated DNA-PAINT and STORM imaging**. a) NanoJ-Fluidics workflow used for STORM and DNA-PAINT imaging. b) Left, full view of a cell showing 5-channel merge of STORM and DNA-PAINT with actin (yellow), vimentin (blue), tubulin (green), clathrin (cyan) and mitochondria (red). Right, zoom of the boxed area. c) Single-channel image of each imaged target (top), with insets (bottom) showing a zoom of the boxed area. A movie corresponding to this experiment is available as Sup. Movie S2. Scale bar corresponds to 10 μm for full images and 2 μm for zoom.

**Fig. 4.**
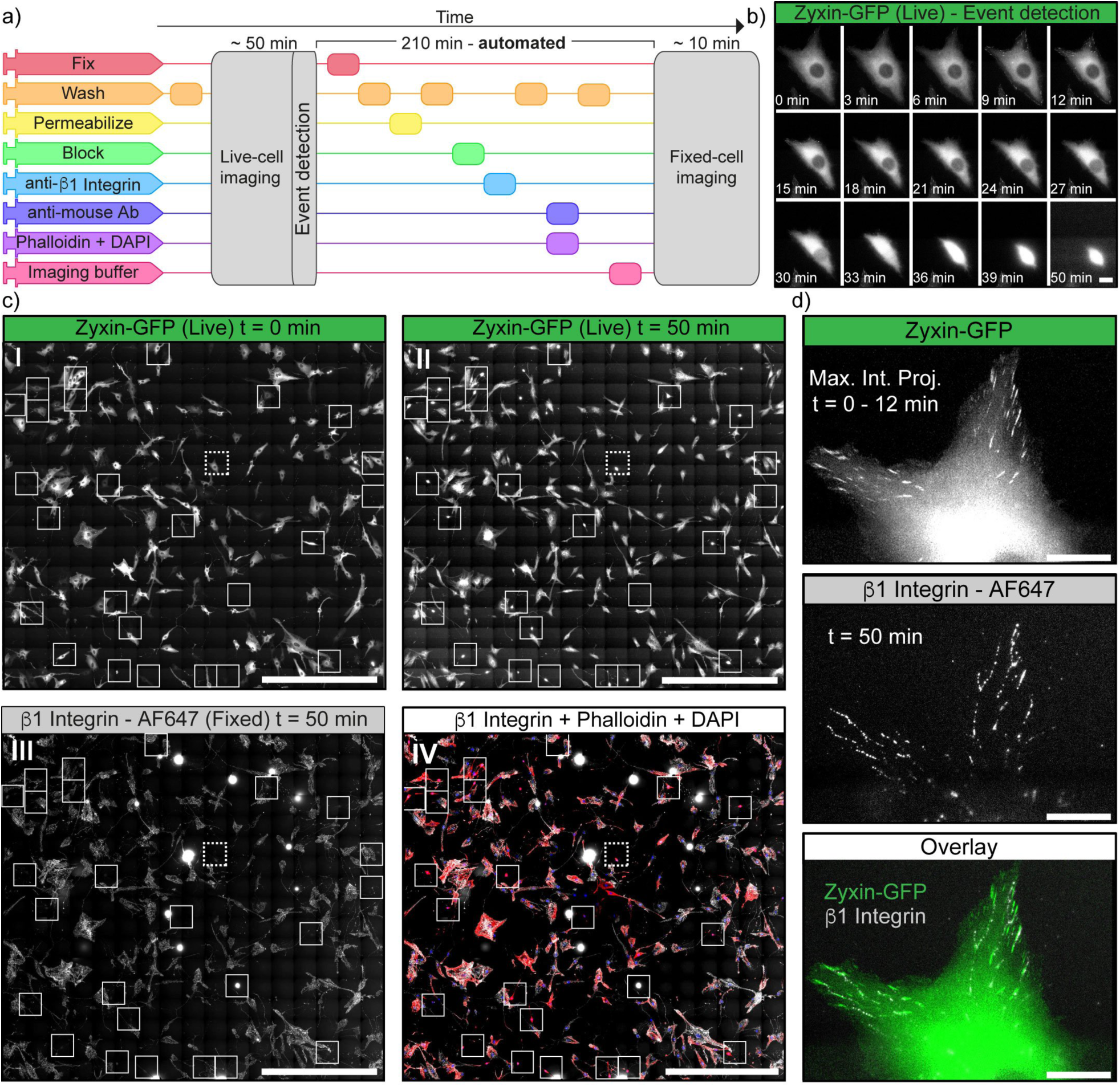
Event-driven fixation of cells upon mitotic rounding. a) NanoJ-Fluidics workflow of the protocol event-driven treatment performed here. b) Stills of RPE1 zyxin-GFP live-cell time-lapse during mitotic rounding. Scale bar, 20 μm.c) Stitched mosaic (17×17 individual regions) of: I-First frame of the live-cell time-lapse; II-Last frame of the live-cell time-lapse; III-RPE1 zyxin-GFP cells immunolabelled for active β1-integrin; IV-Overlay of RPE1 zyxin-GFP cells immunolabelled for active β1-integrin and stained for F-actin (with phalloidin-TRITC) and DNA (with DAPI). Insets represent cells where mitotic rounding was observed (Sup. Movie S3), dashed inset is the cell in b) and d). Scale bar is 1 mm. d) I - Maximum intensity projection of the first 12 min in b); II - Active β1-integrin staining; III - Overlay of both panels. Scale bar, 20 μm.

## Results

**The NanoJ-Fluidics framework**. NanoJ-Fluidics is a complete system, using off-the-shelf components, as well as open-source control software. The hardware component consists of compact LEGO^®^ syringe pumps (Fig. 1a) that can be configured as a multiplexed array of up to 128 syringes (Fig. 1b), plus a peristaltic pump and an open-source Arduino^®^ electronic controller to interface with the microscope acquisition computer (Fig. 1b). By using LEGO parts, the system is cost-effective, robust and repeatable thanks to the high-quality standards and low tolerances of the LEGO manufacturing process. The system performs labelling and treatment protocols that were traditionally done on the bench, but these steps can be directly automated on the microscope stage (Fig. S1). For instance, a wash step will completely remove liquid from the imaging chamber and replenish it with wash buffer. The system is easy to set up (Sup. Note 1), highly modular and compatible with most microscopes and experimental workflows (Fig. S1). NanoJ-Fluidics does not require any design and microfabication process and simply uses common labware (Fig. S2). The software component is provided as an ImageJ/Micro-Manager plugin (10, 11) as well as a stand-alone package for independent fluidics control (Sup. Software). By allowing precise control of each steps in the protocol (Fig. S3), NanoJ-Fluidics provides a highly repeatable and robust way to carry out imaging experiments directly onto the microscope.

**Live-to-fixed super-resolution microscopy**. Although super-resolution microscopy in living cells is achievable with techniques such as SIM (12), RESOLFT (13) and SRRF (14), near molecular scale super-resolution microscopy using single-molecule localisation microscopy (SMLM) is typically limited to fixed-cell imaging. This is due to the requirement for non-physiological conditions, including the use of anti-fading media and high-intensity illumination (15, 16) typically required for commonly used approaches such as STORM(17, 18) and DNA-PAINT (19). NanoJ-Fluidics addresses this issue, by allowing the combination of both dynamic live-cell and subsequent fixed-cell high-resolution observations on the same set of cells.

To demonstrate this, we carried out a sequence of live-cell imaging, fixation, labelling and STORM imaging of actin within COS cells (Fig. 2a). Thanks to its compatibility with most standard microscopes and cell friendly low-illumination requirements, the SRRF approach was chosen to achieve live-cell super-resolution imaging of GFP labelled utrophin calponin homology domain, a validated actin probe (20) (Fig. 2b). SRRF achieved an improved resolution (~175 nm, S4) over the standard wide-field images (HILO, ~270 nm, S4), resolving the assembly and disassembly of actin bundles during cell-shape changes (Fig. 2b and Sup. Movie S1). We then used NanoJ-Fluidics to perform automated fixation, washing, permeabilisation and phalloidin-AF647 staining, a sequence lasting 100 min (Fig. 2a). We finally performed a STORM acquisition on the previously imaged cell after perfusing the optimised imaging buffer (Fig. 2c). The structural detail of the actin organisation observed by STORM is considerably higher than the live-cell observation, with an estimated resolution of ~43 nm. An extended characterisation of the resolutions achieved here is described in Sup. Note 2 and Fig. S4, using NanoJ-SQUIRREL (21). Our system thus provides easy correlative live-cell and fixed-cell super-resolution imaging. Interestingly, these observations also make it possible to evaluate if there are unwanted structural changes to cells caused by the fixation process (shown and discussed in Sup. Note 3).

**Multiplexed super-resolution microscopy with STORM and DNA-PAINT**. Obtaining a super-resolution, high-quality multi-channel image has long been a challenge in SMLM as a consequence of the difficulty to optimise labelling and imaging for many labels simultaneously (22). NanoJ-Fluidics is ideally suited for large channel multiplexing via sequential exchange of fluorescent labels and/or imaging buffers. Furthermore, recent DNA-PAINT modalities can reach more than the typical 2-3 label channels by using antibodies coupled to orthogonal DNA strands, and sequential labelling combined with imaging sequences of PAINT imagers (23–25).

Here, we demonstrate how NanoJ-Fluidics easily handles sequential STORM and DNA-PAINT acquisitions in a simple and optimal manner. Fixed cells were labelled using DNA-coupled primary antibodies targeting mitochondria, vimentin, microtubules and clathrin, as well as fluorescent phalloidin to label actin. NanoJ-Fluidics was set up to carry ++out imaging buffer exchange for STORM imaging of actin (26), followed by two rounds of washing and labelling for 2-colour DNA-PAINT imaging of the other targets (Fig. 3a). The first round imaged mitochondria and vimentin whereas the second round imaged clathrin and tubulin. Fig. 3b-c respectively show the resulting 5-colour image and the individual channels, along with zooms on cellular regions highlighting the high resolution obtained by this combined STORM and DNA-PAINT scheme (~68 nm minimum resolution, except actin rendering which has a 97 nm minimum, characterisation shown in Sup. Note 2 and Fig. S5).

**Fixation-on-event imaging**. NanoJ-Fluidics has also the advantage of allowing sample treatments, such as fixation, at precise times during the experiment. Thanks to the integration of NanoJ-Fluidics with the image acquisition, determining the time of treatment can be triggered by visual queues obtained from the imaging. To demonstrate this capacity, we carried out an experiment observing the state of focal adhesions, as mammalian cells progress into division. Fixation was triggered by the observation of the rounding of the cells as they approach mitosis (27). Also, in order to fully exploit the fluidics automation of NanoJ-Fluidics, we combined it with tiling imaging and image stitching in order to obtain fields-of-view of several millimetres while conserving high-resolution.

We first blocked asynchronous cells in G2 via treatment with a CDK1 inhibitor (RPE1 cells expressing zyxin-GFP). Next, the cell cycle was released by exchanging the inhibitor by growth media using NanoJ-Fluidics (Fig. 4a) and imaged by live-cell time-lapse microscopy (Fig. 4b). When the observed cells reached a minimal area (an indicator of mitotic rounding), the fixative was injected (Fig. 4a-b). A considerable portion of cells (~10%) were found to be fixed in a similar rounding state due to the simultaneous release of the G2 block (insets of Fig. 4c I, Sup. Movie S3). The sample was then immunolabelled for β1-integrin, plus co-stained for F-actin and DNA, then imaged (Fig. 4 c III-VI). Both actin and DNA staining allowed a secondary visual validation of pre-mitotic cell rounding. As we recently described using comparable experiments (28), β1-integrin is shown to retain a similar spatial pattern similar to zyxin when the cells were spread on the substrate, in their pre-division shape (Fig. 4d). We observed that, while zyxin retracts during cell rounding (Fig. 4b), β1-integrin remains in its original position (Fig. 4d and Sup. Movie S3), helping guide daughter cell migration (28).

## Discussion

We introduce NanoJ-Fluidics (Fig. 1) and demonstrate its applicability in multiple experimental contexts: *in-situ* correlative live-to-fixed super-resolution imaging (Fig. 2); multimodal multi-label super-resolution imaging (Fig. 3); and high-content, event-driven correlative live-to-fixed imaging (Fig.4). The reliability provided by the computer-controlled fluidic experiments overcomes the issues of repeatability and low throughput of traditional, non-automated approaches.

Effortless correlative live-cell and fixed-cell super-resolution imaging (Fig. 2) enables a synergy between the dynamic but less spatially resolved information obtained from live-cell imaging and the near molecular-scale structural information provided by SMLM. The NanoJ-Fluidics framework allows to do this *in-situ* without the laborious requirement of relocating multiple cells or performing manual immunolabelling steps. The live-to-fixed transition, and more generally sample treatment, can also be based on specific biological cues, allowing for event-driven correlative live-cell and fixed-cell imaging (Fig. 4). This showcases the broad scope of the NanoJ-Fluidics in multiple and increasingly complex experimental settings, and highlights its potential in combination with recent high-throughput and/or high-content approaches (29–33).

Besides this potential for complex biological interrogation, the simplicity and robustness of the NanoJ-Fluidics facilitates experimental optimisation (Sup. Note 3) (21). To obtain optimal SMLM images, for example, the type, timing and concentration of fixation, permeabilisation and blocking reagents, the antibody concentration, the fluorophores used and the composition of the imaging buffer should be optimised. However, tuning each step individually and especially in combination is a daunting task. The NanoJ-fluidics automation and its integration with imaging allows for a more comprehensive and reliable exploration of the many options available (21, 34–38).

Fluorescent microscopy is widely used for its molecular specificity of labelling and its potential for multi-colour imaging. However, SMLM microscopy approaches tend to be restricted to a couple of colours due to fluorophore photophysics requiring specific imaging buffers. A flexible and easy-to-use fluidics system, such as NanoJ-Fluidics is a significant advance in this context: the easy combination of STORM (22) with DNA-PAINT (23–25, 39), as we demonstrate in Fig. 3, and the potential to perform sequential labelling protocols (e.g. bleaching or antibody removal (40, 41)) can extend high spatial resolution multiplexing far beyond what live-cell fluorescent microscopy can currently achieve.

NanoJ-Fluidics is easy (and fun) to set up and use. Its highly modular design means that the system can be easily adapted for specific experiments, such as by detaching syringe pump modules to be kept at different temperature (Fig. S1). There are several advantages to the NanoJ-Fluidics design compared to standard flow cells: glass-bottom chambers are easier to prepare in comparison to microfluidic chips; unwanted air bubbles formed in the tubing are easily dissipated before corrupting the sample; being a non-pressurised system means that there is higher tolerance to flow-rate fluctuations. Notwithstanding, flow cells have the general advantage of requiring smaller volumes of reagents. Automated pipetting robots are also a good alternative, being particularly efficient in multiwell high-content experiments (42), but are generally restricted to highly specialised laboratories. The open-source software component of the NanoJ-Fluidics provides an Application Programming Interface (API) and plugin framework that allows third party syringe pumps, such as those commercially available, to be controlled by the common NanoJ-Fluidics graphical user interface (GUI, Fig. S3). This means that the automation protocols established here are directly applicable to pre-existing or alternative hardware.

In this work we have chosen to focus mainly on the application of NanoJ-Fluidics to live-cell imaging and SMLM approaches, as each SMLM imaging requires a finely-tuned chemical environments to produce high-resolution images. However, our fluidic exchange system can be used in any imaging experiments that would benefit from an automated exchange of the sample media. Our Micro-Manager plugin already allows for NanoJ-Fluidics to be fully integrated into microscopy acquisition software, enabling a seamless combination between the imaging and fluid exchange protocol. In conclusion, NanoJ-Fluidics makes high-content multi-modal microscopy experiments tractable and available to a large audience of researchers, while improving reliability, optimisation and repeatability of imaging protocols.

**Software and Hardware Availability**. NanoJ-Fluidics follows open-source software and hardware standards, it is part of the NanoJ project (14, 21, 43). The steps to assemble a complete functioning system are described in https://github.com/HenriquesLab/NanoJ-Fluidics/wiki.

## ACKNOWLEDGEMENTS

We thank Prof. Ralf Jungmann at Max Planck Institute (MPI) of Biochemistry Munich for reagents and advice. This work was funded by grants from the UK Biotechnology and Biological Sciences Research Council (BB/M022374/1; BB/P027431/1; BB/R000697/1) (R.H., P.M.P. and R.F.L.), the UK Medical Research Council (MR/K015826/1) (R.H.), the Wellcome Trust (203276/Z/16/Z) (S.C. and R.H.) and the Centre Nationnal de la Recherche Scientifique (CNRS ATIP-AVENIR program AO2016) (C.L.). PA. was supported by a PhD fellowship from the UK’s Biotechnology and Biological Sciences Research Council. C.L.D. was supported by PhD funding from the Medical Research Council, UK (1214605). Research by B.B. was supported by UCL, Cancer Research UK (C1529/A17343), and MRC (MC_CF12266).

## AUTHOR CONTRIBUTIONS

PA. and R.H. devised the hardware and wrote the software. P.A., P.M.P., C.L. and R.H. planned experiments. Experimental data sets were acquired by P.M.P. (Fig. 2), G.C, F.B.R. and C.L. (Fig. 3), P.A. and C.L.D. (Fig. 4). Data was analysed by P.A., P.M.P. and S.C. while G.C., B.B., C.L. and R.H. provided research advice. The paper was written by P.A. P.M.P., R.F.L., C.L. and R.H. with editing contributions of all the authors.

## Methods

**Cell lines**. COS7 cells were cultured in phenol-red free DMEM (Gibco) supplemented with 2 mM GlutaMAX (Gibco), 50 U/ml penicillin, 50 μg/ml streptomycin (Pen-strep, Gibco) and 10% fetal bovine serum (FBS; Gibco). hTERT-RPE1 cells stably expressing Zyxin-GFP (1) were cultured in DMEM F-12 Glutamax (Gibco), with 10% FBS, 3.4% sodium bicarbonate (Gibco), 1% Penstrep. All cells were grown at 37°C in a 5% CO_2_ humidified incubator. Cell lines have not been authenticated.

**Plasmids**. GFP-UtrCH was a gift from William Bement (2) (Addgene plasmid #26737).

**Antibody conjugation**. Secondary antibodies (see bellow for details) were labelled with DNA strands (see bellow for details) as previously described (3). In short, secondary antibodies were concentrated via amicon 100 kDa spin filters to 2-6 mg/ml. 50-100 μg of antibody was labelled using a Maleimide-Peg2-succinimidyl ester for 90 min at 40x molar excess at 4°C on a shaker. Crosslinker stocks of 10 mg/ml in DMF were diluted in 1x PBS to reach 40x molar excess in 5 μl, which were subsequently added to the antibody. After the reaction had been done, unreacted crosslinker was removed via a zeba spin column. Thiolated DNA was reduced using DTT for 2h at room temperature. DTT was separated from the reduced DNA via a Nap5 column and fractions containing DNA were concentrated via 3 kDa amicon spin filters. The reduced DNA was then added to the antibody bearing a functional maleimide group in 25x molar excess and incubated over night at 4°C on a shaker in the dark. Antibody-DNA constructs were finally purified via 100 kDa amicon spin filters. DNA-PAINT labelling: - For Mitochondria: Goat anti-Mouse (A28174, ThermoFisher) with I1 (docking: 5’-TTATACATCTA-3’; imager: 5’-CTAGATGTAT-ATTO655-3’); - For Vimentin: Goat anti-chicken (Abcam, ab7113) with I2 (docking: 5’-TTAATTGAGTA-3’; imager: 5’-GTACTCAATT-Cy3B-3’); - For Clathrin: Goat anti-Rabbit (A27033, ThermoFisher). Goat anti-Chicken (ab7113, Abcam) with I3 (docking: 5’- TTTCTTCATTA-3’; imager: 5’-GTAATGAAGA-Cy3B-3’); - For alpha-tubulin: Donkey anti-Rat (A18747, ThermoFisher) with I4 (docking: 5’-TTTATTAAGCT-3’; imager: 5’-CAGCTTAATA-ATTO655-3’). Sequences for DNA-PAINT strands were obtained from (4). Both thiolated and fluorophore conjugated DNA strands were obtained from Metabion.

**NanoJ-Fluidics framework**. We provide the detailed instructions to easily build and use the system in a regular biology lab, as well as the software enabling its control and automation (Sup. Note 1).

**Live-to-Fixed Super-Resolution imaging**. The NanoJ-Fluidics syringe pump array was installed on a Nikon N-STORM microscope equipped with 405, 488, 561 and 647 nm lasers (20, 50, 50 and 125 mW at the optical fiber output). One individual syringe pump module containing the fixative was kept within the incubator of the microscope at 37°C. All steps after cell transfection were performed on the microscope, using NanoJ-Fluidics. COS7 cells were seeded on ultraclean (5) 25 mm diameter thickness 1.5H coverslips (Marienfeld) at a density of 0.3−0.9×10^5^ cells/cm^2^. One day after splitting, cells were transfected with a plasmid encoding the calponin homology domain of utrophin fused to GFP (GFP-UtrCH) using Lipofectamin 2000 (Thermo Fisher Scientific) according to the manufacturer’s recommendations. Cells were imaged 1-2 days post transfection in culture medium using an Attofluor cell chamber (Thermofisher), covered with the lid of a 35 mm dish (Thermofisher), that was kept in place using black non-reflective aluminum tape (T205-1.0—AT205, THORLABs).

Cells were fixed at 37°C for 15 minutes with 4% paraformaldehyde in cytoskeleton-preserving buffer (1X PEM, 80 mM PIPES pH 6.8, 5 mM EGTA, 2 mM MgCl2) (6). After fixation cells were permeabilised (1X PEM with 0.25% Triton-X) for 20 min, blocked with blocking buffer (5% Bovine Serum Albumin (BSA) in 1X PEM) for 30 minutes, and stained with Phalloidin-AF647 (Molecular Probes, 4 units/mL) for 30 minutes.

Laser-illumination Highly Inclined and Laminated Optical sheet (HILO) imaging of Utrophin-GFP in live COS7 cells was performed at 37 °C and 5% CO_2_ on a Nikon N-STORM microscope. A 100x TIRF objective (Plan-APOCHROMAT 100x/1.49 Oil, Nikon) with additional 1.5x magnification was used to collect fluorescence onto an EMCCD camera (iXon Ultra 897, Andor), yielding a pixel size of 107 nm. For timelapse imaging, 100 raw frames (33 ms exposure) were acquired once every 10 minutes (with the illumination shutter closed between acquisitions) for 150 minutes with 488 nm laser illumination at 4% of maximum output. STORM HILO imaging of Alexa Fluor 647-phalloidin in fixed cells was performed on the same system. 50,000 frames were acquired with 33 ms exposure and 642 nm laser illumination at maximum output power with 405 nm pumping when required. STORM imaging was performed in GLOX buffer (150mM Tris, pH 8, 1% glycerol, 1% glucose, 10mM NaCl, 1% β-mercaptoethanol, 0.5 mg/ml glucose oxidase, 40 μg/ml catalase) supplemented with Phalloidin-Alexa Fluor 647 (1 U/mL).

**Multiplexed Super-Resolution microscopy with Ex-change-PAINT and STORM.** COS-7 cells (obtained from ATCC) were seeded on 18mm, 1.5H glass coverslips (Menzel-Gläser). 24 hours after seeding, they were fixed using 4% PFA, 4% sucrose in PEM buffer at 37°C (6). After blocking in phosphate buffer with 0.022% gelatin, 0.1% Triton-X100 for 1.5 hours, cells were incubated with primary antibodies overnight at 4°C: mouse monoclonal anti-TOM20 (BD Bioscience # 612278), rabbit polyclonal anti-clathrin heavy chain (abcam ab21679), chicken polyclonal anti-vimentin (BioLegend # 919101) and rat antialpha-tubulin (mix of clone YL1/2 abcam # 6160 and clone YOL1/34 Millipore CBL270). After rinses, they were incubated with Exchange-PAINT secondary antibodies coupled to DNA sequences: goat anti-mouse I1, goat anti-chicken I2, goat anti-rabbit I3 and donkey anti-rat I4 for 1.5 hours at RT (see Ab conjugation section for antibody and sequence details). After rinses, they were incubated in phalloidin-Atto488 (Sigma) at 12.5 μM for 90 min at RT and imaged within a few days. For STORM/PAINT imaging, the NanoJ-Fluidics array was installed on an N-STORM microscope (Nikon) equipped with 405, 488, 561 and 647 nm lasers (25, 80, 80 and 125 mW at the optical fiber output). First, a STORM image of phalloidin-ATTO488 was performed in buffer C (PBS 0.1M pH7.2, 500 mM NaCl) using 30,000 frames at 30 ms/frame at 50% power of the 488 nm laser. After injection of the I1-ATTO655 and I2-CY3B imagers in buffer C, 60,000 frames were aquired in an alternating way (647 nm at 60% and 561 nm at 30%) to image TOM20 and vimentin, respectively. After three rinses with buffer C, I3-Cy3B and I4-ATTO655 were in buffer C were injected, and 60,000 frames were aquired in an alternating way (561 nm at 30% 647 nm at 60%) to image clathrin and microtubules, respectively. All imager strands were used at concentration between 0.25 and 2 nM.

**Event Detection and Live-to-Fix Imaging**. hTERT-RPE1 cells stably expressing Zyxin-GFP were incubated with 9 mM Ro-3306 (Enzolife Sciences ALX-270-463) to inhibit CDK1 activity for 15-20 hours. Inhibition was released by replacing drug containing media by fresh media at the microscope immediately before imaging. Cells were imaged using a Nikon Eclipse Ti microscope (Nikon) equipped with an Neo-Zyla sCMOS camera (Andor), LED illumination (CoolLED) and a 60X objective (Plan Apo 60X/1.4 Oil, Nikon). Images were acquired every 3 min until enough cells had underwent mitotic rounding. At that point, 16% warmed PFA was added to cells in media to a final concentration of 4%, and incubated at room temperature for 20 minutes. They were then washed 3 times and 0.2% Triton was added for 5 minutes. 5% BSA in 1X PBS was used to block for 30 min at room temperature, before activated β1 Integrin (Abcam #ab30394) primary antibody was added. After incubation and washing, Phalloidin-TRITC (Sigma-Aldrich) and anti-mouse AF647 antibody (Invitrogen) were added. All of these steps were performed automatically using the NanoJ-Fluidics platform.

**SMLM and SRRF image reconstruction**. For Fig. 2 images were reconstructed using NanoJ-SRRF (7) (TRPPM for live cell data and TRM for fixed cell data with a magnification of 4). Drift was estimated using the inbuilt function in NanoJ-SRRF and correction applied during SRRF analysis. For figure 3 localizations were detected using the N-STORM software (Nikon), and exported as a text file before being filtered and rendered using ThunderSTORM (8). Chromatic aberration between the red (561 nm) and far-red (647 nm) channels were corrected within the N-STORM software using polynomial warping, and remaining translational drift between acquisition passes were aligned manually on high-resolution reconstructions.

FRC values were obtained using NanoJ-SQUIRREL after reconstruction of original data separated into two two different stacks composed of odd or even images (9). NanoJ-SRRF, NanoJ-SQUIRREL and ThunderSTORM are available in Fiji (10).

---

**Fig. S1.**
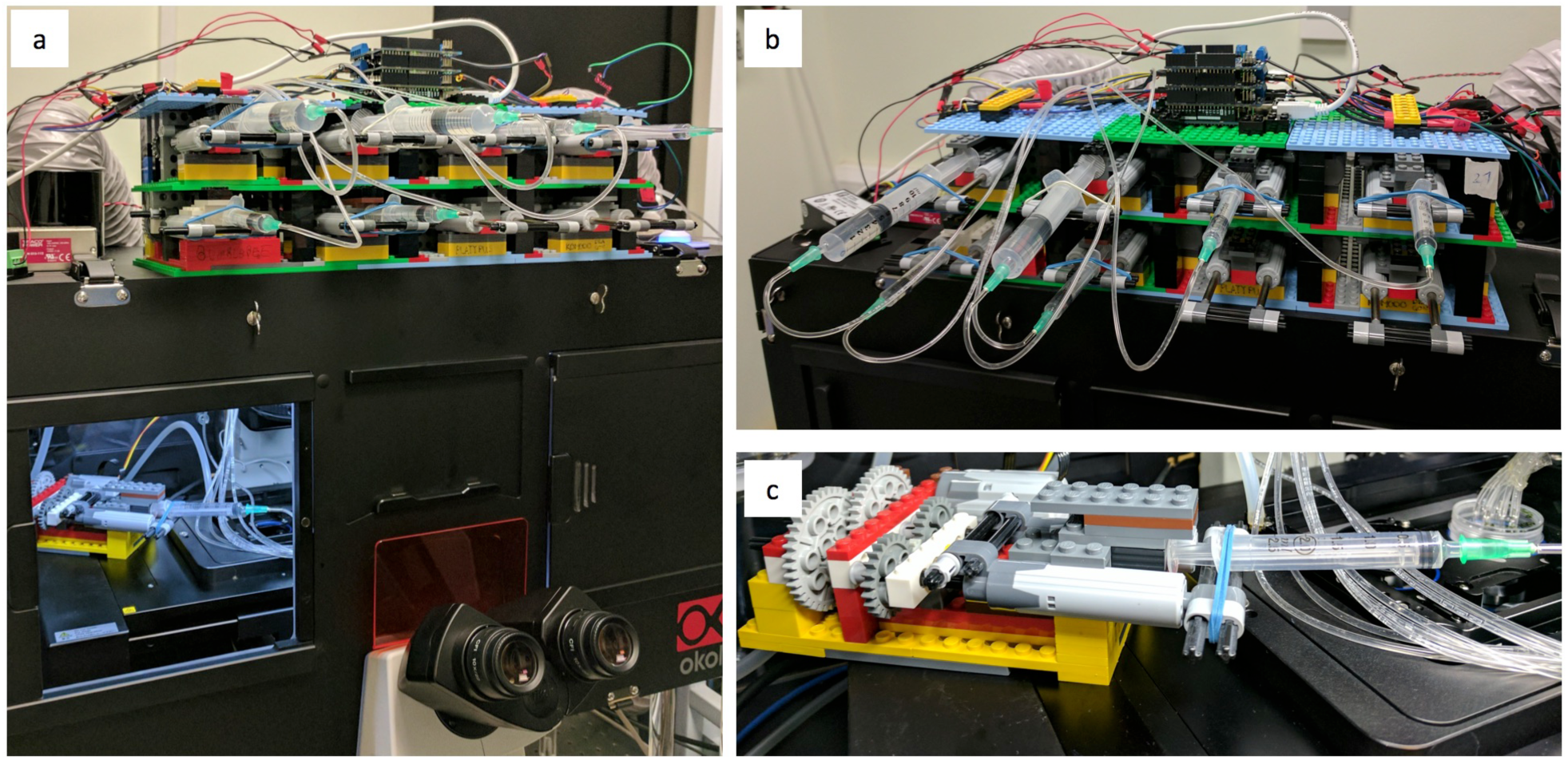
Assembled NanoJ-Fluidics system on a microscope. a) View of assembled pump array with syringes loaded on top of a Nikon N-STORM microscope, an individual syringe pump unit sits inside the incubator and is kept at 37°C in the microscope incubator, allowing the use of reagents equilibrated at the same temperature as the sample such as fixatives. b) Top view of the syringe pump array. c) Zoom into the individual syringe pump unit inside incubator.

**Fig. S2.**
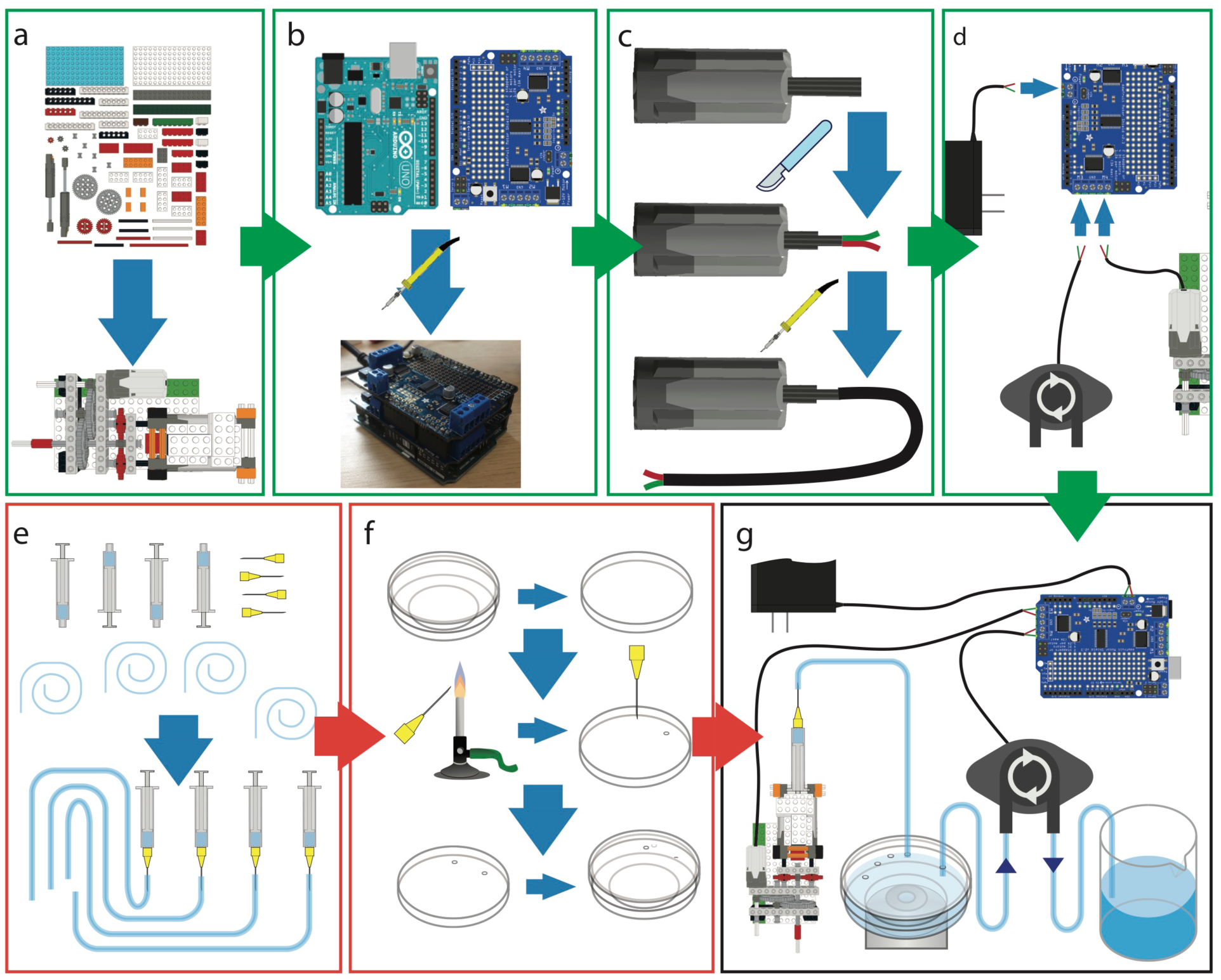
NanoJ-Fluidics pump assembly. There are 4 steps to assemble a NanoJ-Fluidics system (a-d), and 2 steps to prepare an experiment (e-f). a) Build the syringe pumps from LEGO bricks. b) Assemble the electronic controller. c) Prepare the motor cables for wiring. d) Connect the pumps and power supply to the controller. e) Prepare the syringes. f) Prepare lid of the cell culture dish. g) Thread syringes on the dish lid and mount syringes on LEGO pumps.

**Design and capabilities**. The NanoJ-Fluidics system is designed primarily for the sequential exchange of liquids in a glass-bottom cell culture dish (or equivalent supports, such as ATTOfluor coverlslip holders for example). It consists of several parts: LEGO based syringe pumps responsible for delivering reagents to the sample (Fig. 1a); electronics responsible for controlling the pumps (Fig. 1b-d); a peristaltic pump responsible for removing reagents from the sample (Fig. 1d); fluid handling disposable components (Fig. 1e-f). The syringe pumps are designed to accommodate syringes of any size, to be cost-effective and simple to assemble. A syringe pump unit uses a simple gear and actuator system to translate fast motor motion to smooth and slower linear motion, to enable consistent fluid flow. The result is that the flow-rate will depend on the gears, motor speed and the syringe’s internal diameter. The pumps can be calibrated by measuring the flow-rate obtained at a given motor speed and syringe volume. Given these reference values, the flow-rate for any other syringe diameter (*R_d_*) can be determined using Eq. S1:

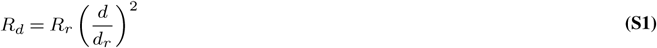

Where *d* is the syringe diameter, *d_r_* is the reference diameter and *R_r_* is the reference rate. With the current design, a 1mm BD Plastipak syringe with an internal diameter of 4.699mm was measured to obtain a nominal flow-rate of 2.3 ± 0.08 μL/s at maximum motor speed. Based on this value, Table S1 summarizes the limits for different BD Plastipak syringe volumes calculated from Eq. S1. Table S1 also shows the maximum loading volume for each syringe given that this will be limited by the length of the linear actuators on the pumps.

**Table S1.** Flow-rates obtained from Eq. S1 for several BD Plastipak syringes. Minimal flow rate is limited by the minimal rotation speed achievable by the DC LEGO motors without stalling.

| Nominal Volume (mL) | Max. Volume (mL) | Inner Diameter (mm) | Max. Flow Rate ($\mu\text{L/s}$) | Min. Flow Rate ($\mu\text{L/s}$) |
| --- | --- | --- | --- | --- |
| 1 | 0.75 | 4.699 | 2.3 | 0.6 |
| 2 | 2.5 | 8.7 | 7.9 | 2.0 |
| 5 | 4.8 | 11.989 | 15.0 | 3.7 |
| 10 | 6.5 | 14.427 | 21.7 | 5.4 |
| 20 | 11 | 19.05 | 37.8 | 9.5 |
| 50 | 22 | 26.594 | 73.7 | 18.4 |

Both the syringe pump array (for media injection) and peristaltic pump (for media removal) are digitally run using an Arduino UNO micro-controller and Adafruit Motorshield digital-to-analogue motor control. The Motorshield is an additional electronic board that connects to the top of the Arduino controller, enabling it to run up to 4 LEGO motors. An Arduino can be stacked with up to 32 Motorshields, limiting the controller to a maximum of 128 pumps, which should be sufficient for most NanoJ-Fluidics applications. We provide custom open-source firmware that enables the Arduino-based electronics to be programmatically controlled by a connected computer. We also provide a Java-based graphical user interface (GUI) for simple control of the fluidics sequences (Fig. S2).

The software interface can be set to automatically perform a sequence of liquid injection and liquid removal steps (Sup. Note Fig. S3) allowing an entire sample fixation or labelling protocol to be carried as shown in Fig. 2-3, 4.

**Assembly**. The steps to prepare a complete functioning NanoJ-Fluidics system (Fig. S2 and S1) consist of assembling the pump array, controller, and wiring them together (Fig. S2a-d). Once the array and controller have been assembled, each experiment only requires the lab-ware component to be prepared anew (Fig. S2 e-g). A full step-by-step guide for assembly and preparation can be found in https://github.com/HenriquesLab/NanoJ-Fluidics/wiki.

NanoJ-Fluidics can also be used with more traditional microfluidics devices, such as Polydimethylsiloxane (PDMS) chips, however we have focused on providing a series of protocols using off-the-shelf labware (e.g. glass-bottom cell culture dishes). This approach makes the framework more accessible to laboratories that do not otherwise have the facilities to produce PDMS devices. When using culture dishes the primary use of the system is limited to liquid exchange, this effectively reproduces many laboratory protocols including cell fixation, immunolabeling, drug treatment, among others.

**Fig. S3.**
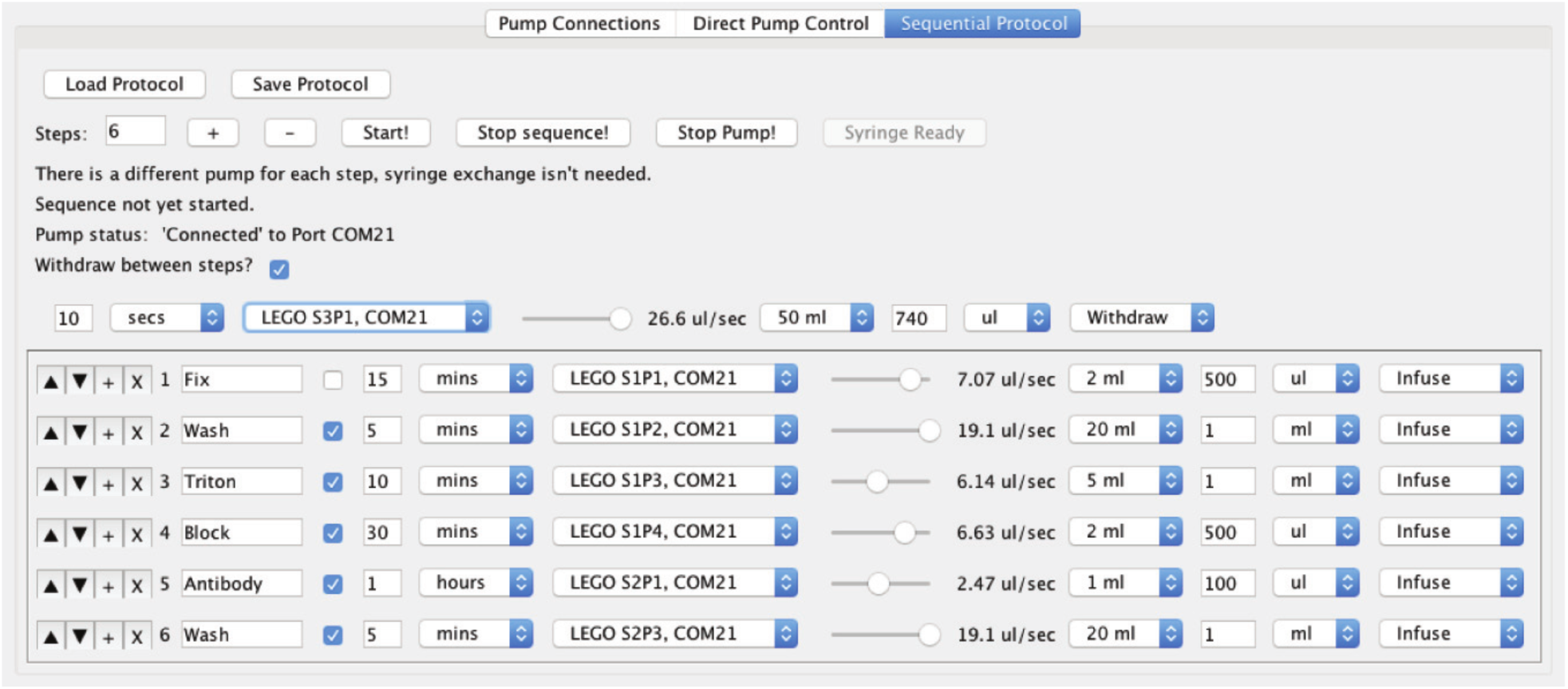
User Interface. Screenshot of the NanoJ-Fluidics sequential control user interface.

**Software interface and experiment automation**. Control of the NanoJ-Fluidics pump array can be achieved in one of three ways: by using the provided Java-based GUI (Fig. S3); RS232 commands to the Arduino board; or by using the Application Programming Interface (API). A description of these is available at https://github.com/HenriquesLab/NanoJ-Fluidics/wiki.

The RS232 command scheme enables researchers with programming experience to run the controller directly in their own software packages. However, the GUI was designed to enable any user to directly run any number of Arduino controllers and pumps, as well as design a sequence of steps associated to an experimental protocol (Fig. S3).

NanoJ-Fluidics is controlled by a set of Java classes described in the API. For example, these can be used to enable a protocol generated in the GUI to be programmatically started and manipulated by outside applications, such as a microscope control software such as Micro-Manager (1). This enables complex automated protocols, such as pre-screening in live imaging to try and capture specific events, such as mitosis (2, 3), before fixing and further analysis the sample (Fig. 4). Therefore, by using the NanoJ-Fluidics GUI and API, any combination of sample acquisition and liquid manipulation protocols can be performed in a completely automated manner.

## Supplementary Note 2: Resolution mapping

**Fig. S4.**
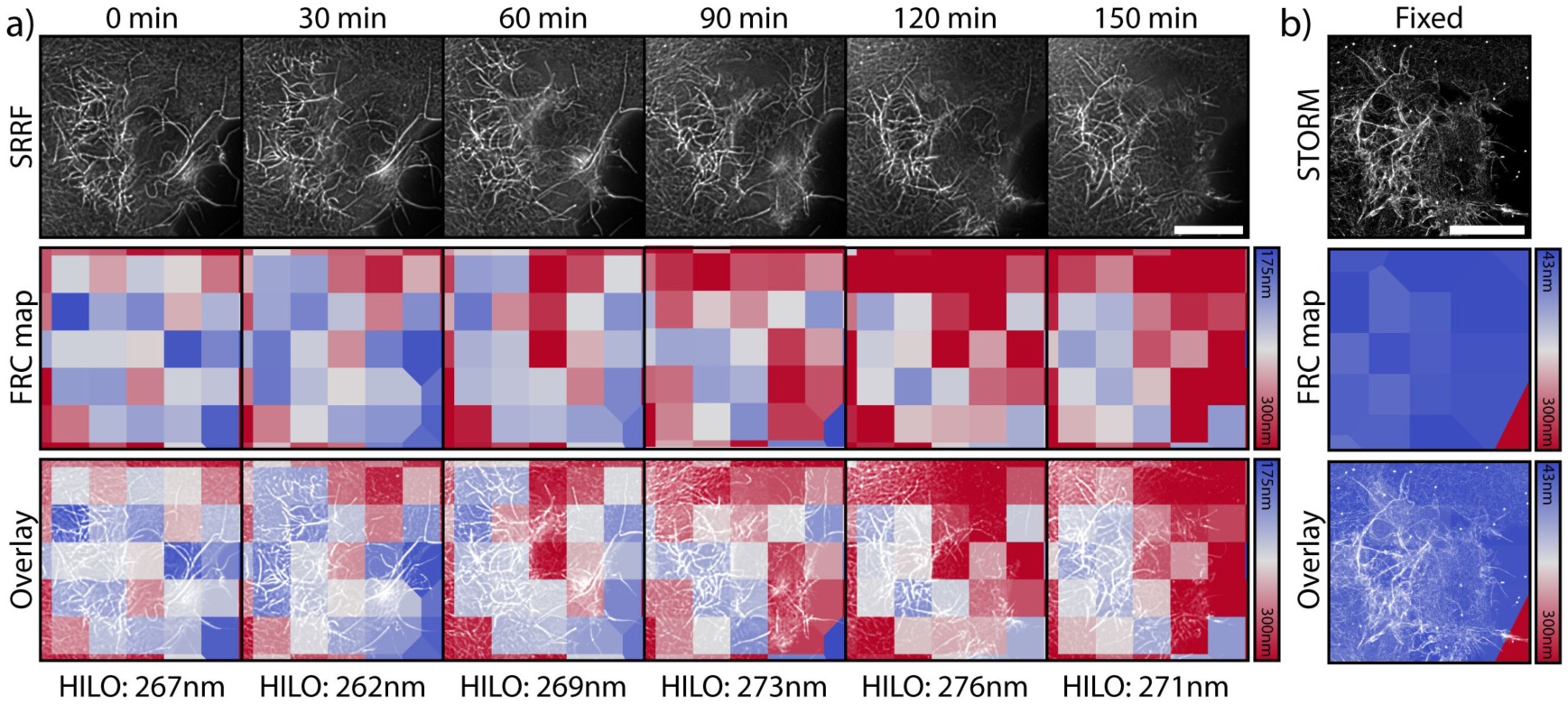
Fourier Ring Correlation (FRC) resolution mapping for Fig. 2 and S1 using NanoJ-SQUIRREL. a) Individual live-cell SRRF frames at different time-points pre-fixation (top-row), equivalent FRC map (middle-row) and overlay between SRRF frames and the corresponding FRC map (lower-row); bottom values show average resolution calculated for the diffraction limited equivalent images. The mean HILO resolutions for each image are shown beneath. b) Individual STORM rendering acquired post-fixation (top); equivalent FRC map (middle); overlay between STORM frame and FRC map. Resolution maps calculated through NanoJ-SQUIRREL (4). All scale bars are 10 μm.

**Fig. S5.**
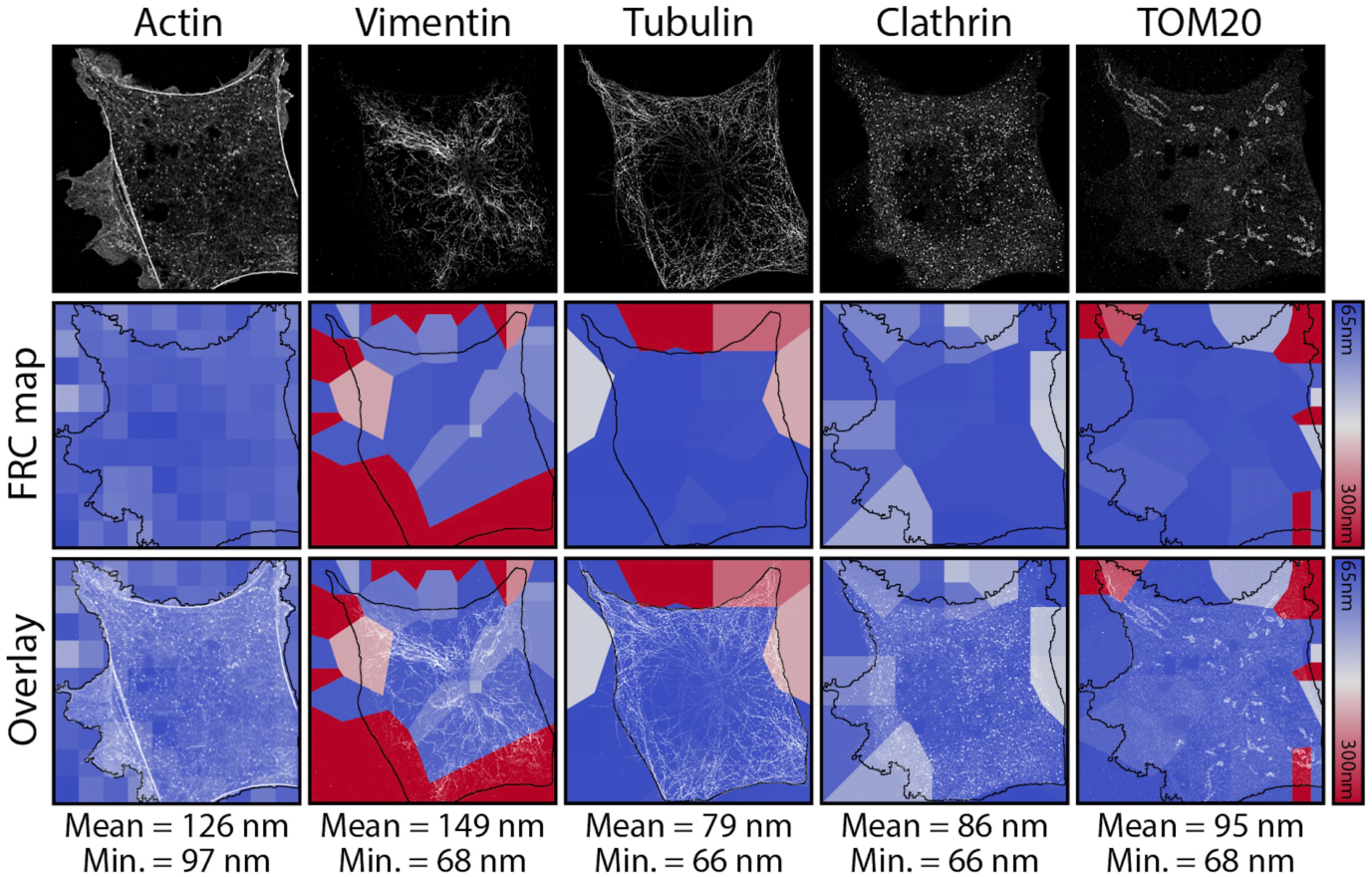
Fourier Ring Correlation (FRC) resolution mapping for Fig. 3 and S2 using NanoJ-SQUIRREL. Individual DNA-PAINT super-resolution renderings (top-row), equivalent FRC map (middle-row) and overlay between SRRF frames and the corresponding FRC map (lower-row). Resolution maps calculated through NanoJ-SQUIRREL. Black outlines in middle and bottom rows indicate the cell shape mask used for calculation of mean and minimum resolutions across each channel.

To estimate the local resolution achieved in the super-resolution renderings associated to the main figures, we carried out analysis using the Fourier Ring Correlation method (5), recently modified in (4) to generate a resolution map. Fig. S4 shows that the best resolution achieved for live-cell imaging with SRRF is 175nm, and 43nm for fixed-cell imaging with STORM. Fig. S5 shows that the best resolution achieved in the DNA-PAINT channels is ~67nm, with the exception of the actin channel which has an estimated best resolution of 97nm.

## Supplementary Note 3: Nanoscale morphological changes between pre- and post-fixation

**Fig. S6.**
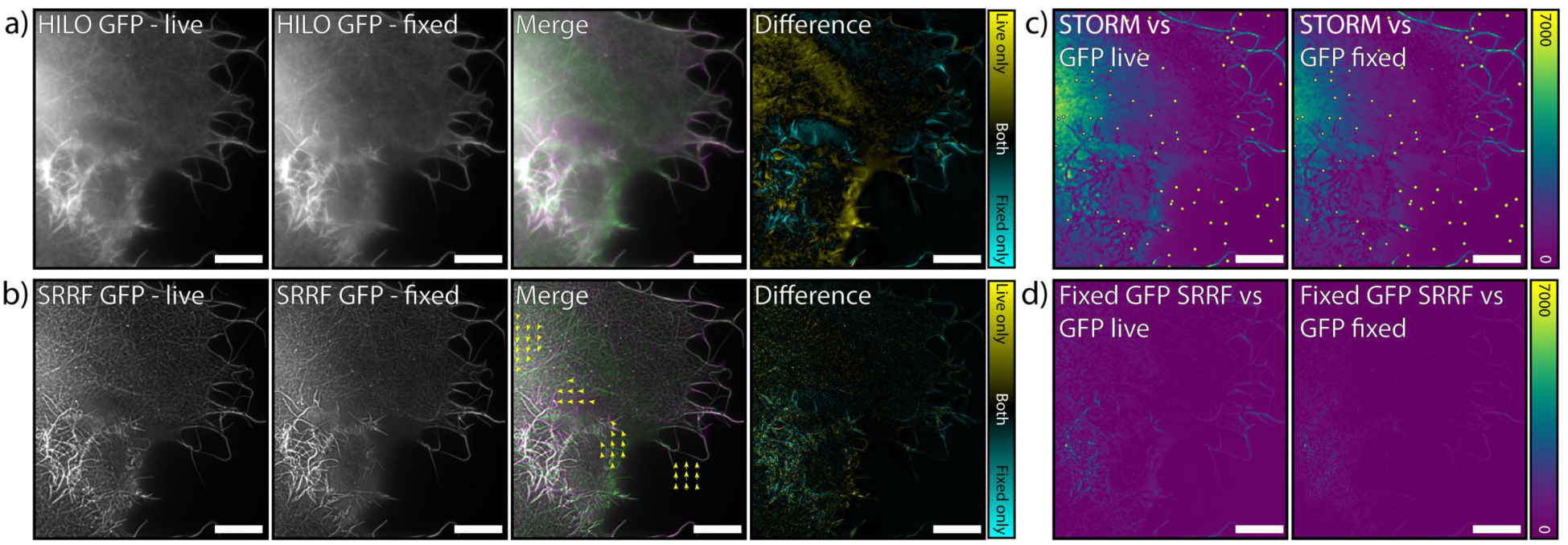
Analysis of changes in cell morphology and labelling pre- and post-fixation. a) Comparison of the distribution of GFP-labelled Utrophin in the cell shown in Fig. 2 and Movie S1, imaged in HILO. The last timepoint pre-fixation (‘HILO GFP - live’) and an image post-fixation of the same region (‘HILO GFP - fixed’) are shown. ‘Merge’ shows an overlay of the live (green) and fixed (magenta) images. ‘Difference’ shows the result of subtracting the fixed image from the live image. b) As in (a), except using the SRRF reconstructions of the data. The yellow arrows in ‘Merge’ show parts of the cell which moved during fixation as measured using the elastic channel registration tool in NanoJ-SQUIRREL (4). c) Error maps generated between the (fixed) STORM reconstruction of actin in this cell when using the live-cell GFP HILO image (left) or the fixed-cell GFP HILO image (right) as a reference. d) Error maps generated between the SRRF reconstruction of GFP in the fixed cell when using the live-cell GFP HILO image (left) or the fixed-cell GFP HILO image (right) as the reference. Scale bars = 10 μm.

To verify that there were no major alterations in cell morphology during fixation, we compared images of the same cell immediately before and after fixation. Fig. S6a shows pre- and post-fixation images for HILO imaging of the cell shown in Fig. 2, and Fig. S6b shows these changes for SRRF reconstructions of the same region. Overlaying both the SRRF and HILO images pre- and post-fixation shows that the morphology of the cell remains largely constant during fixation. There is movement of the bright filaments within the cell body and filopodia at the cell periphery on a sub-micron scale. This degree of movement is comparable to or smaller than the frame-to-frame movement shown in Movie S1. There is also a loss of fluorescence intensity during fixation in the top left portion of the region shown in Fig. S6.

**Movie S1.**
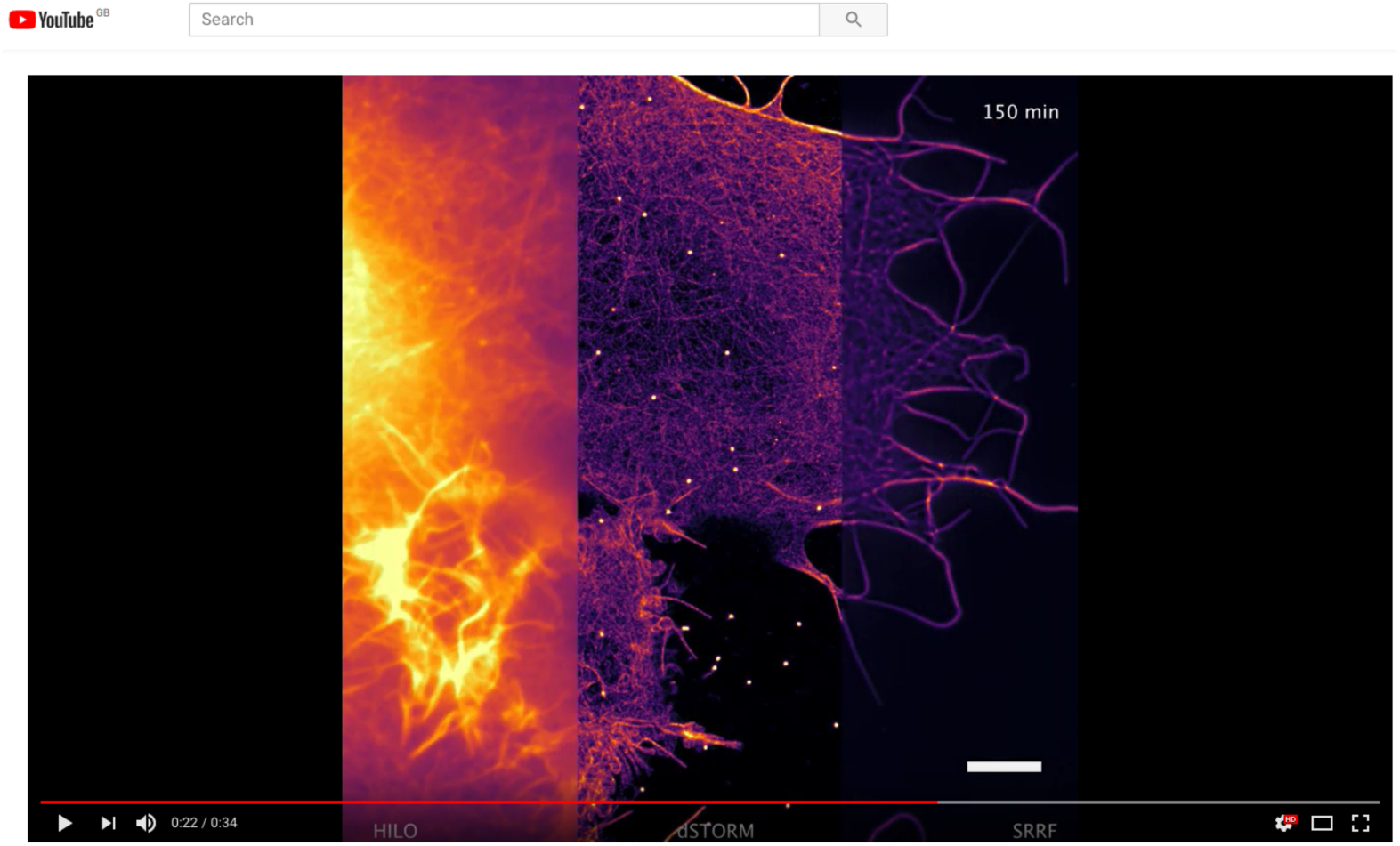
Live-to-Fix Super-Resolution Imaging with NanoJ-Fluidics. Movie corresponding to Fig. 2. HILO and HILO-SRRF live imaging of transiently transfected COS7 cells expressing UtrCH-GFP. After live-imaging the same cells were fixed and stained with phalloidin-AF647 for STORM imaging. All steps were performed using the NanoJ-Fluidics syringe pump array without taking the sample from the microscope. Scale bars correspond to 10 μm. Movie can be found in: https://youtu.be/v6yfqVgKbDE

**Movie S2.**
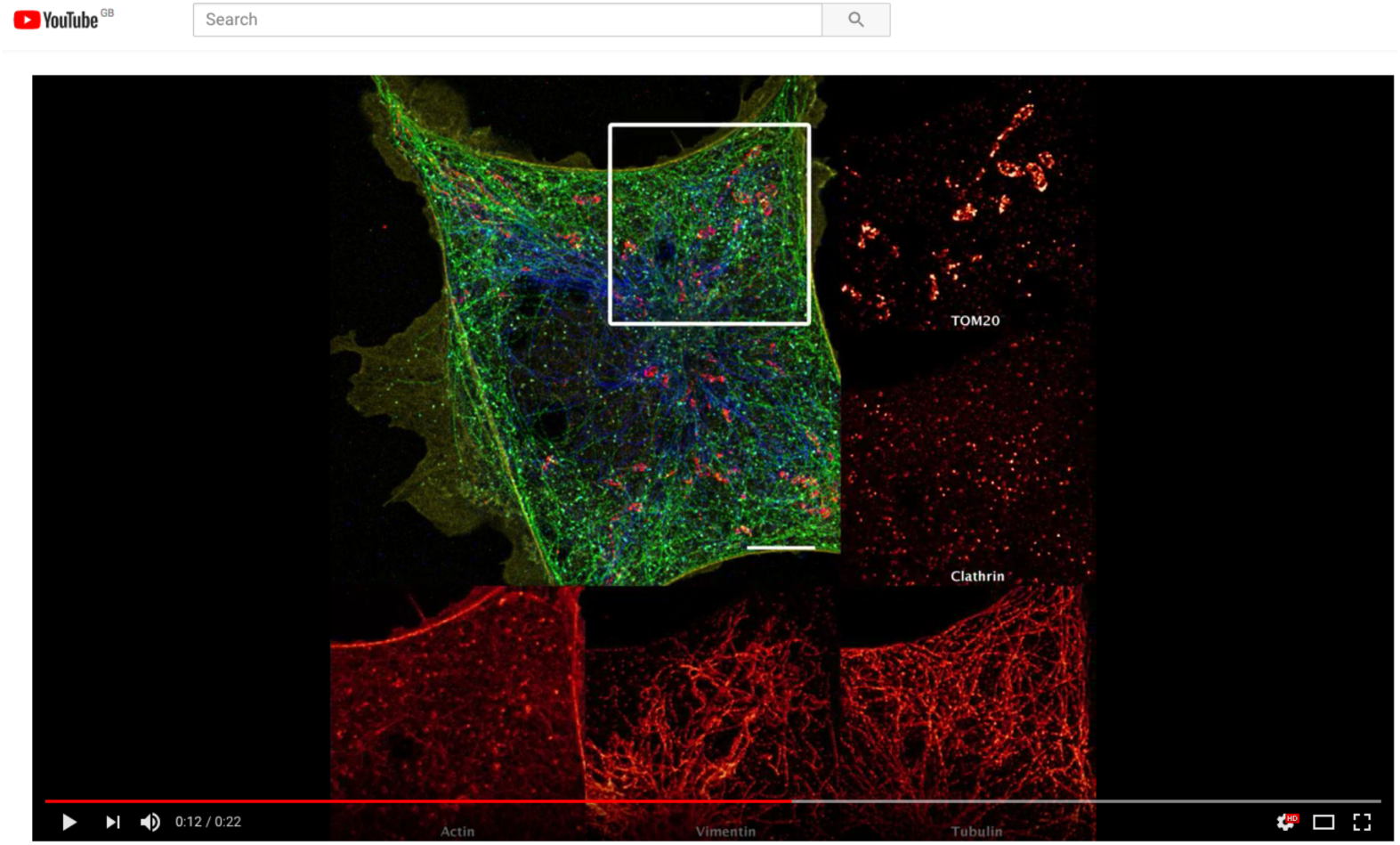
Automated Multiplex Super-Resolution with NanoJ-Fluidics. Movie corresponding to Fig. 3. STORM imaging of COS7 cells stained with phalloidin-Atto488, and DNA-PAINT imaging of vimentin, TOM20, tubulin and clathrin the same cell. Imaging buffer exchange steps were performed using the NanoJ-Fluidics syringe pump array directly on the microscope stage. Scale bars corresponds to 10 μm. Movie can be found in: https://youtu.be/VMV1iQOYYhY

## Supplementary Movie 3: Event Detection and Live-to-Fix Imaging with NanoJ-Fluidics

**Movie S3.**
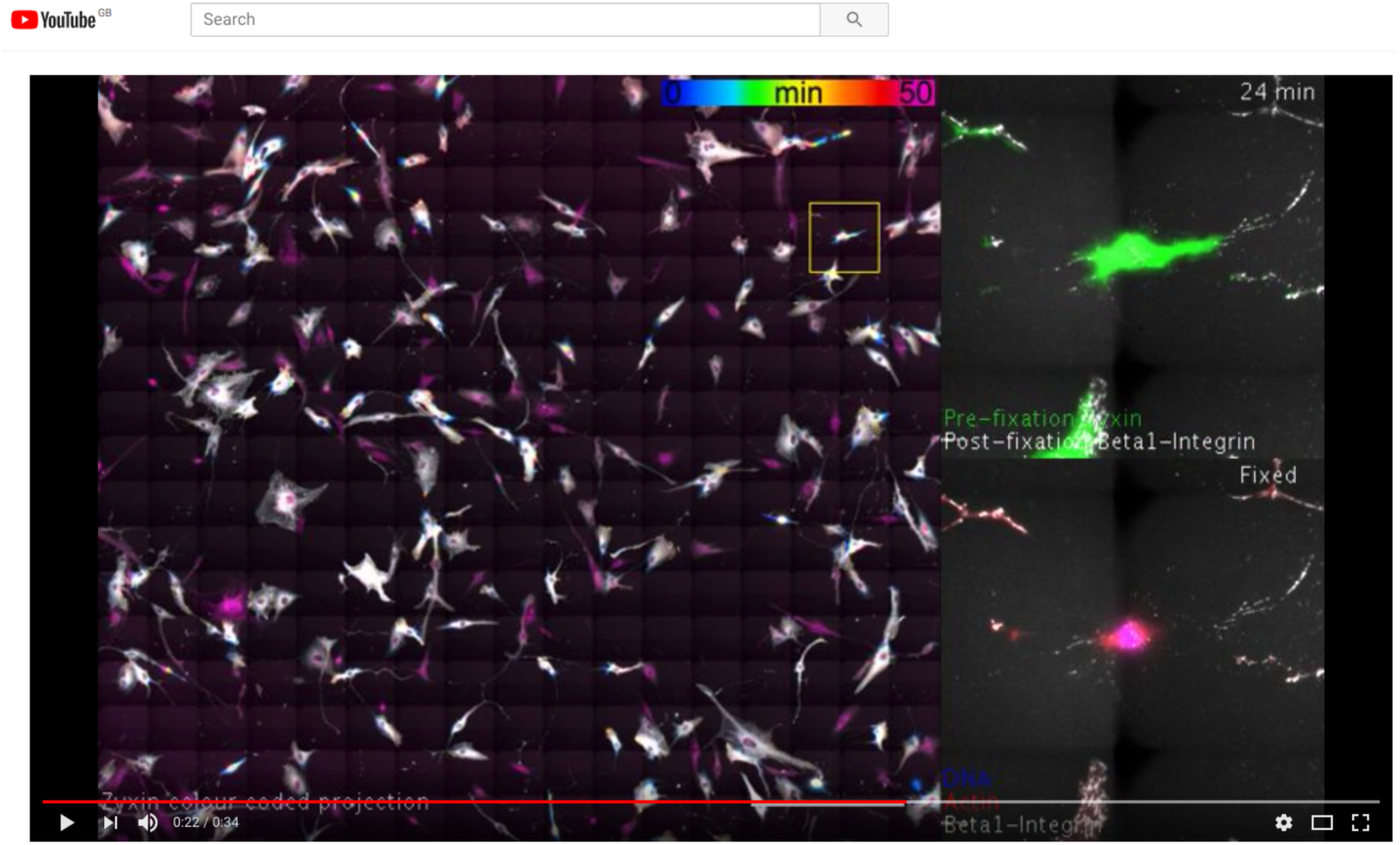
Event-driven live-to-fixed imaging with NanoJ-Fluidics. Movie corresponding to Fig. 4. Left: colour coded time projection of a stitched mosaic (17×17 individual regions), following RPE1 cells stably expressing zyxin-GFP undergoing mitotic rounding. Upon enough cells rounded (event detection) cells were fixed and immunolabelled for active β1-integrin and stained for actin and DNA. On the top right corner, overlay of zyxin-GFP in live cells with active β1-integrin after fixation. First and last regions correspond to cells not undergoing mitotic rounding, whereas all remaining regions display RPE1 cells rounding and showcase the colocalization of active β1-integrin (post-fixation staining) with focal adhesions (zyxin-GFP). Bottom right corner, overlay of active β1-integrin, phalloidin and DAPI staining after fixation. Scale bar corresponds to 0.5 mm. Movie can be found in: https://youtu.be/sbeP1AvKWDA

